## Supplementary material for "Barrier properties of Nup98 FG phases ruled by FG motif identity and inter-FG spacer length": Supp

Contents:

**Supplementary Figures 1-2** with legend

**Supplementary Tables 1-3**

**Supplementary Note 1:** Complete amino acid sequences of all engineered FG domain variants and reference wild-type FG domains

**Supplemental References**

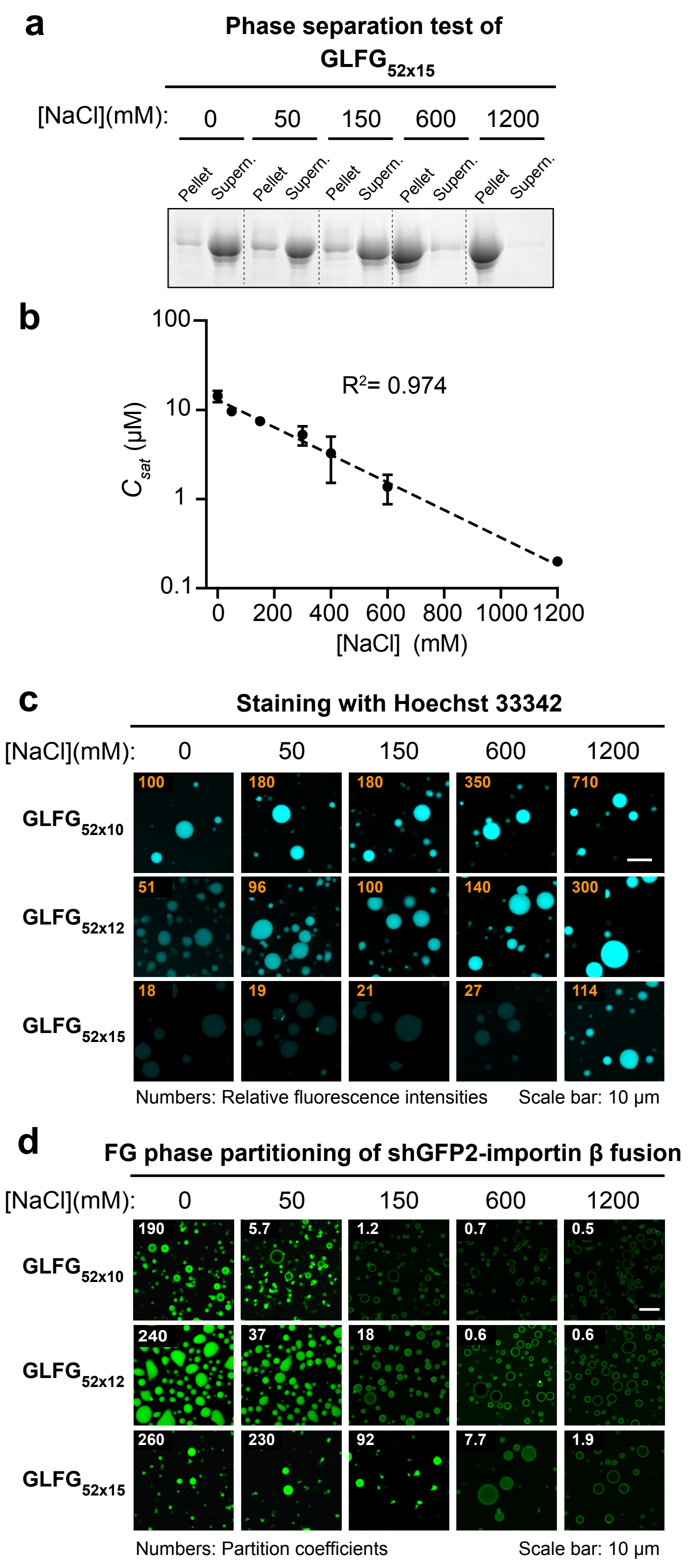


**Supplementary Figure 1: Increasing salt concentration results in hypercohesive strict FG phases.**

**(a)** Phase separation of $GLFG_{52x15}$ was analysed at a concentration of 10 µM in buffers containing 50 mM Tris/HCl pH 7.5, 5 mM DTT + indicated NaCl concentrations. The assay was performed three times on independent samples with similar results, and a representative image is shown. A full scan of the gel with molecular weight markers is provided in the Source Data file.

**(b)** *Saturation concentration* ($C_{sat}$) for phase separation of $GLFG_{52x15}$ is plotted against [NaCl] in the assay buffers. Measurements were performed three times with independent samples, and mean values are shown with S.D. as error bars. Phase separation tests of $GLFG_{52x15}$ at [NaCl] = 0 and 50 mM were repeated at [$GLFG_{52x15}$] = 20 µM for determination of saturation concentrations. The mean values were fitted to a simple exponential function (dashed line) with the R-squared value indicated. Note that higher NaCl concentrations lead to stronger cohesive interactions.

**(c)** FG phases assembled from the indicated variants were stained with Hoechst 33342 in buffers containing 50 mM Tris/HCl, 5 mM DTT + indicated concentrations of NaCl. The numbers in orange indicate the fluorescence intensities of the Hoechst dye inside the FG phases. The fluorescence intensities are relative to that of $GLFG_{52x12}$ at 150 mM NaCl (arbitrarily set to 100). Note that elevated intensities correlate with hyper-cohesive interactions caused by high [NaCl].

**(d)** FG phases assembled from the indicated variants were challenged with an shGFP2-Importin β fusion (shGFP2 is an engineered FG-phobic moiety) at the indicated concentrations of NaCl. Scanning settings/ image brightness were adjusted individually due to the large range of signals. The numbers in white refer to the partition coefficients of the shGFP2-Importin β fusion into the FG phases (fluorescence ratios in the central regions of the particles to that in the surrounding buffer). The use of shGFP2-Importin β fusion ruled out that the GFP dissociates from Importin β in varying spacer lengths or under varying salt concentrations. Note that hyper-cohesive interactions, caused by high [NaCl], impede the entry of shGFP2-Importin β into the GLFG phases for all three variants. **(c & d)** Each of the assays was performed twice on independent samples with similar results, and representative images are shown.

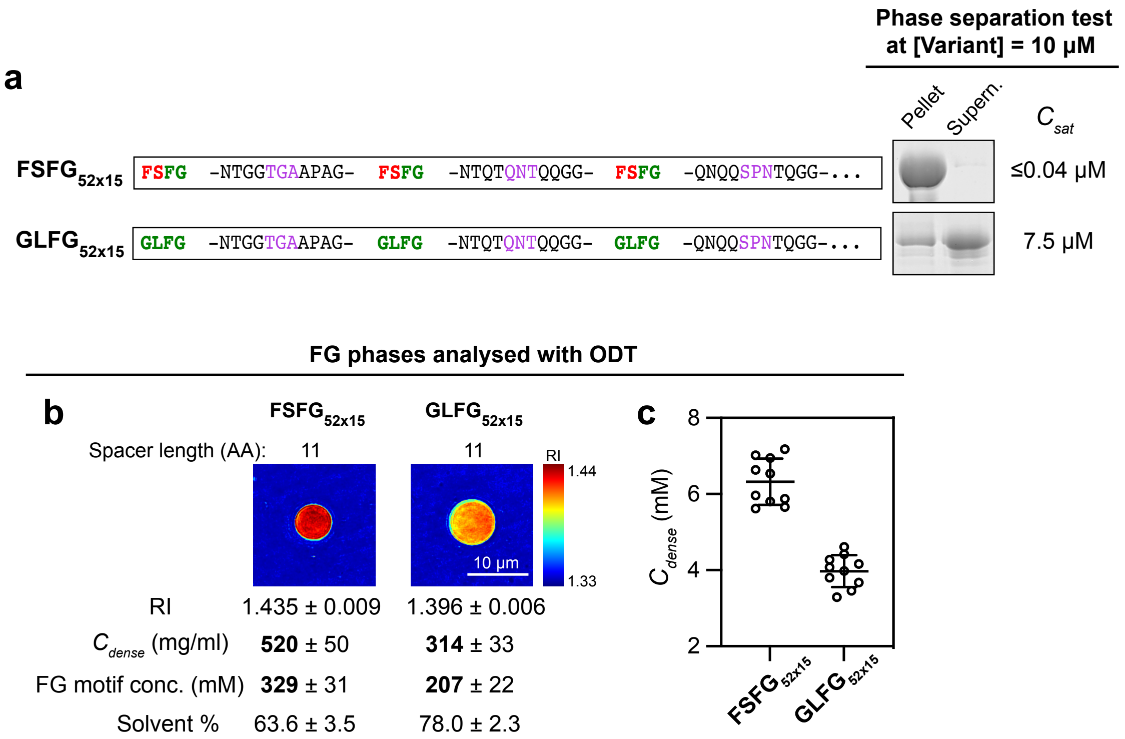


**Supplementary Figure 2: FSFG motifs lead to stronger cohesive interactions and more negative ΔG for phase separation than the GLFG motifs.**
**(a)** Phase separation of $FSFG_{52x15}$ and $GLFG_{52x15}$ was analysed at a concentration of 10 µM. Samples of the obtained pellets (FG phase) and supernatants were loaded for SDS-PAGE at equal ratio (6%), followed by Coomassie blue-staining for quantification. Saturation concentration, $C_{sat}$, for phase separation of each is taken as the concentration of the supernatant. $C_{sat}$ of $FSFG_{52x15}$ is so low that it could not be quantified accurately and thus the upper limit is given. Note that the FSFG motifs lead to a stronger phase separation propensity than the GLFG motifs as they lead to a much lower $C_{sat}$.
**(b & c)** Phases assembled from $FSFG_{52x15}$ and $GLFG_{52x15}$ were analysed by optical diffraction tomography (ODT) as described in the main text to determine the refractive indices (RI) and protein concentrations within the condensed phases ($C_{dense}$). For each, ten independent FG particles were analysed. Experiments described in **(a)** and **(b-c)** are two independent experiments. Representative phase maps and mean values ± S.D. are shown in **(b)**. Individual data points in molar concentration are plotted with the bars representing mean ± S.D. in **(c)**. Source data of this figure are provided in the Source Data file. Note that according to Eq.1, ΔG for phase separation of $FSFG_{52x15}$ is ≤ -29.7 kJ/mol, and thus ΔG for $FSFG_{52x15}$ and $GLFG_{52x15}$ differ by at least 14.1 kJ/mol.

| | *Hs*Nup98 | *Bf*Nup98 | *Dm*Nup98 | *Ce*Nup98 | *Dd*Nup220 | *Sc*Nup116 | *Tt*Mac98A | **GLFG$_{52x12}$** |
|---|---|---|---|---|---|---|---|---|
| Number of residues in FG domain | 443 | 427 | 528 | 442 | 669 | 680 | 622 | 623 |
| Number of FG motifs | 39 | 40 | 46 | 36 | 56 | 47 | 42 | 52 |
| Number of FG motifs per 100 AA | 8.8 | 9.4 | 8.7 | 8.1 | 8.4 | 6.9 | 6.8 | 8.3 |
| Number of occurrences of sub-types of FG motif per 100 AA (% of all FG motifs in each domain): | | | | | | | | |
| GLFG | **1.8 (20.5)** | **3.0 (32.5)** | 1.1 (13.0) | **2.5 (30.6)** | **3.4 (41.1)** | **2.9 (42.6)** | **4.3 (64.3)** | **8.3 (100)** |
| GIFG | 0.0 (0.0) | 0.0 (0.0) | 0.0 (0.0) | 0.0 (0.0) | 0.0 (0.0) | 0.29 (4.3) | 0.32 (4.8) | 0.0 (0.0) |
| SLFG | 0.45 (5.1) | 0.47 (5.0) | 0.95(10.9) | **2.5 (30.6)** | 1.2 (14.3) | 0.29 (4.3) | 0.16 (2.4) | 0.0 (0.0) |
| SIFG | 0.23 (2.6) | 0.23 (2.5) | 0.0 (0.0) | 0.90(11.1) | 0.0 (0.0) | 0.29 (4.3) | 0.0 (0.0) | 0.0 (0.0) |
| AFG | 1.1 (12.8) | 0.70 (7.5) | **2.1 (23.9)** | 0.45 (5.6) | 0.15 (1.8) | 1.0 (14.9) | 0.0 (0.0) | 0.0 (0.0) |
| PFG | 0.90(10.3) | 0.70 (7.5) | 0.57 (6.5) | 0.23 (2.8) | 1.6 (19.6) | 0.59 (8.5) | 0.16 (2.4) | 0.0 (0.0) |
| FSFG | 0.23 (2.6) | 0.23 (2.5) | 0.0 (0.0) | 0.23 (2.8) | 0.0 (0.0) | 0.0 (0.0) | 0.0 (0.0) | 0.0 (0.0) |
| Other FG motifs | 4.1 (46.2) | 4.0 (42.5) | 4.0 (45.7) | 1.4(16.7) | 1.9 (23.2) | 1.5 (21.3) | 1.8 (26.2) | 0.0 (0.0) |
| Occurrences of FG-like motifs per 100 AA: | | | | | | | | |
| GLFA | 0.0 | 0.47 | 0.0 | 0.0 | 0.0 | 0.0 | 0.0 | 0.0 |
| GLFS | 0.23 | 0.0 | 0.0 | 0.0 | 0.0 | 0.15 | 0.0 | 0.0 |
| GFLG | 0.0 | 0.0 | 0.0 | 0.0 | 0.0 | 0.0 | 0.16 | 0.0 |
| LG | 1.1 | 1.6 | 0.57 | 0.0 | 0.60 | 0.0 | 1.1 | 0.0 |
| IG | 0.45 | 0.23 | 0.0 | 0.0 | 0.0 | 0.0 | 0.16 | 0.0 |
| Occurrences of other hydrophobic residues per 100 AA: | | | | | | | | |
| F | 1.8 | 0.23 | 1.9 | 0.23 | 0.0 | 1.0 | 0.64 | 0.0 |
| Y | 0.45 | 0.0 | 0.0 | 0.0 | 0.0 | 0.0 | 0.0 | 0.0 |
| W | 0.0 | 0.0 | 0.0 | 0.0 | 0.0 | 0.0 | 0.0 | 0.0 |
| L+I+V+M | 3.6 | 0.94 | 2.8 | 1.6 | 1.8 | 4.1 | 3.1 | 0.0 |
| Total number of carbons in the sidechains of F,Y,W,L,I,V,M per 100 AA: | | | | | | | | |
| | 111 | 100 | 99 | 89 | 86 | 87 | 88 | 92 |

**Supplementary Table 1: Occurrences of different types of FG motifs and hydrophobic residues in wild-type Nup98 FG domains from indicated eukaryotic species, in comparison to a sequence-regularized FG domain variant.** Nup98 FG domain homologs from a wide range of species: *Homo sapiens* (representing vertebrates), *Branchiostoma floridae* (representing lancelets), *Drosophila melanogaster* (representing insects), *Caenorhabditis elegans* (representing nematodes), *Dictyostelium discoideum* Nup220 (representing amoebas), *Saccharomyces cerevisiae* Nup116 (representing fungi), and MacNup98A from *Tetrahymena thermophila* (representing ciliates) were analysed. The most common type(s) of FG motif among these FG domains are in bold. In counting of all, the GLEBS domains (~40-50 residues) were omitted. Note that the overall hydrophobicity of vertebrate Nup98 FG domains is higher than the others. This may correlate with the fact that these FG domains are typically glycosylated.

| | Mac98A FG domain | $GLFG_{52x10}$ | $GLFG_{52x11}$ | $GLFG_{52x12}$ (standard) | $GLFG_{52x13}$ | $GLFG_{52x14}$ | $GLFG_{52x15}$ |
|---|---|---|---|---|---|---|---|
| Total amino acids (AA) | 622 | 520 | 572 | 623 | 676 | 728 | 780 |
| **GLFG motifs** | 27 | **52** | **52** | **52** | **52** | **52** | **52** |
| Other FG motifs | 15 | **0** | **0** | **0** | **0** | **0** | **0** |
| FG motifs per 100 AA | 6.8 | 10 | 9.1 | 8.3 | 7.7 | 7.1 | 6.7 |
| FG-like motifs | 14 | **0** | **0** | **0** | **0** | **0** | **0** |
| Amino acid frequencies in inter-GLFG spacers (%) | | | | | | | |
| A | 12.8 | 17.5 | 15.6 | 15.8 | 16.0 | 15.2 | 15.4 |
| C | 0.0 | 0.0 | 0.0 | 0.0 | 0.0 | 0.0 | 0.0 |
| D | 0.0 | 0.0 | 0.0 | 0.0 | 0.0 | 0.0 | 0.0 |
| E | 0.0 | 0.0 | 0.0 | 0.0 | 0.0 | 0.0 | 0.0 |
| **F** | 3.9 | **0.0** | **0.0** | **0.0** | **0.0** | **0.0** | **0.0** |
| G | 30.7 | 18.2 | 26.3 | 26.1 | 26.0 | 27.6 | 27.4 |
| H | 0.0 | 0.0 | 0.0 | 0.0 | 0.0 | 0.0 | 0.0 |
| **I** | 0.8 | **0.0** | **0.0** | **0.0** | **0.0** | **0.0** | **0.0** |
| K | 0.4 | 1.0 | 0.8 | 0.7 | 0.6 | 0.6 | 0.5 |
| **L** | 3.1 | **0.0** | **0.0** | **0.0** | **0.0** | **0.0** | **0.0** |
| **M** | 1.8 | **0.0** | **0.0** | **0.0** | **0.0** | **0.0** | **0.0** |
| N | 12.6 | 17.5 | 15.6 | 15.6 | 15.6 | 15.2 | 15.2 |
| P | 5.8 | 7.6 | 7.1 | 7.2 | 7.2 | 7.1 | 7.2 |
| Q | 10.1 | 14.0 | 12.3 | 12.5 | 12.6 | 12.3 | 12.4 |
| R | 0.2 | 0.0 | 0.0 | 0.0 | 0.0 | 0.0 | 0.0 |
| S | 1.8 | 1.9 | 2.5 | 2.4 | 2.3 | 2.3 | 2.3 |
| T | 15.8 | 22.3 | 19.7 | 19.7 | 19.6 | 19.8 | 19.7 |
| **V** | 0.2 | **0.0** | **0.0** | **0.0** | **0.0** | **0.0** | **0.0** |
| **W** | 0.0 | 0.0 | 0.0 | 0.0 | 0.0 | 0.0 | 0.0 |
| **Y** | 0.0 | 0.0 | 0.0 | 0.0 | 0.0 | 0.0 | 0.0 |

**Supplementary Table 2: Number of FG- or FG-like motifs and amino acid frequencies (%) in inter-GLFG spacers of the wild-type MacNup98A (Mac98A) FG domain and variants.** All sequences contain a *Tetrahymena thermophila* GLEBS domain (44 residues) which was omitted in all counting. Note that the spacers of the variants do not contain any hydrophobic residues (F, Y, W, L, I, V nor M, in bold).

| **Protein name** | **Plasmid** | **Encoding for** | **Reference** |
|---|---|---|---|
| mCherry | pSF779 | $His_{14}$-TEV-mCherry-Cys | [1] |
| EGFP | pSF1526 | $His_{14}$-MBP- *bd*SUMO-mEGFP | [2] |
| efGFP_8Q | pDG2936 | $His_{14}$-*bd*SUMO-efGFP_8Q | [2] |
| efGFP_8R | pSF2892 | $His_{14}$-*bd*SUMO-efGFP_8R | [2] |
| NTF2 | pDG2121 | rat NTF2 | [2] |
| *hs*RanGDP | pDG2961 | $His_{14}$-ZZ-*sc*SUMO-Cys-*hs*Ran | this study |
| *hs*Transportin | pKK006 | $His_{10}$-mEGFP-TEV-*hs*Transportin | [1] |
| *hs*M9-EGFP | pSNG136 | $His_{14}$-*bd*SUMO-*hs*M9-mEGFP | this study |
| *sc*Importin β (*sc*Impβ) | pMR676 | $His_{14}$-*bd*SUMO-*sc*Kap95p | [1] |
| *sc*IBB-EGFP | pSF807 | $His_{14}$-TEV-*sc*Srp1p$_{2\text{-}63}$-mEGFP | [1] |
| *hs*Importin β (*hs*Impβ)* | pDG2305 | $His_{14}$-MBP-*bd*SUMO-*hs*Imp_beta | [2] |
| *hs*IBB-sffrGFP7$^{NTR}$ * | pDG2899 | $His_{14}$-*bd*SUMO-*hs*IBB-sffrGFP7 | [2] |
| *hs*Xpo1/*hs*CRM1 | pTG-A42 | $His_{10}$-ZZ-TEV-*hs*CRM1 | [3] |
| *hs*RanQ69L$_{1\text{-}180}$ | pTG-A418 | $His_{10}$-ZZ-TEV-*hs*RanQ69L $_{1\text{-}180}$ | [3] |
| NES-EGFP | pTG-A450 | $His_{14}$-TEV-PKI01-mEGFP-Cys | [4] |
| GFP$^{NTR}$_3B7C | pDG2779 | $His_{14}$-*bd*SUMO-GFP$^{NTR}$3B7C | [2] |
| sffrGFP4$^{NTR}$ | pDG2805 | $His_{14}$-*bd*SUMO-sffrGFP4 | [2] |
| sffrGFP4 25R→K | pSF2885 | $His_{14}$-*bd*SUMO-sffrGFP4 complete R-K mutant | 2 |
| TetraGFP$^{NTR}$ | pDG2913 | $His_{14}$-*bd*SUMO-GFP$^{NTR}$3B7C M225F | this study |
| shGFP2-Impβ | pSF2051 | $His_{14}$-*bd*SUMO-shGFP2-*sc*Kap95p | [5] |

**Supplementary Table 3:** Proteins used as permeation probes and corresponding bacterial expression constructs in this study. Plasmid numbers are unique identifiers. For each a His-tag-free (cleaved via SUMO/ TEV/ NEDD8 sites) version was used. * human Importin β was used for forming complex with *hs*IBB-sffrGFP7$^{NTR}$. Otherwise, yeast (*sc*) Importin β was used for forming complexes with molecules containing the *sc*IBB.

### Supplementary Note 1:
#### Complete amino acid sequences of engineered FG domain variants and reference wild-type FG domains

*Coloured in red: GLEBS domain (44 residues)

**Wild-type *Tetrahymena thermophila* macronuclear Nup98A (Mac98A) FG domain**
**Plasmid: pHBS418**

MFGNTGGGGLFGNTQTQQTGGGLFGQPQQTQFGQTGATGGGLFGGATNTFGGGGGGLFGGNNNQQTNPTAGGGIFGQGT
TGLGGAPAQTGGGLFGAPQNNQGGGLFGGGTTTGGGMFGNQANTQTGGGGLFGGPSQPTTQPPAFSLNNPTTGGGGLFGQ
PANTMGGNNGGLFGGQTNSFGANNNMLGNNNRPQGAGIFGGATTTAPTGNTGMFGGIGANNGGGGLFGMNNTNTNPTGGF
GATNPTAGGGGLFGGGATTTGGGGLFGGGNTQGGGLLGTANTTAGGLLGGGFNMNNNTGGILGQTNNQFGLGSFGTNNNA
AAAPFQPKASANGVLTKPNEKNLCYAISNGTDFCIFELALTQRKLVKAGQLKPGAQQAGGMFGQPAQGGNGLFGGGGAAT
TTPFGGAQNGNLFGGQNTQAQGGGLFGAPVNNAATGAGGGLFGAKPAATTTGGGLFGQMPAQTGGFLGNTATQPAGGGLF
GGATTTQAPGGGGGGGLFGGNTTAATTGGGLFGGNTQTGGATGGLFGGQQPNNQGGLFLNTGNANNANTGGGLFGGATTT
PATGGGLFGGSTNTQPGLATGGGLFGNNQGASQPAAQGGLFGGAAPQQNSLFGGATAGGQTGGLFGGATGATQQQGGGLF
GQTASNPTQGGGLFGAANPGLGGAAA

**GLFG**$_{52x12}$
**Plasmid: pSNG036**

GLFGNTGGAPAGGLFGNTQTQQGGGLFGQPQQTQGGGLFGQTGATTGGGLFGGATNTAPGGLFGGGGGNPTGGLFGGNNN
QQTGGLFGQGTTQTGGGLFGAPQNNQGGGLFGGGTTTTGGGLFGANTQTGGGGLFGGPSQPTTAGLFGSNNPTTGGGLFG
QPANTNNGGLFGGQTNNQASGLFGANNQPPTNGLFGNNNKPQTAGLFGGATTTGNTGLFGGANNTGGGGLFGNNTNNPTG
GLFGATNPAGGGGLFGGGATTGGGGLFGGGNTQTGGGLFGTANTTTAGGLFGGGNTQPQNGLFGNNNTPATGGLFGQTNN
AAPQGLFGGTNNNAASGLFGQKPASANGVLTKPNEKNLCYAISNGTDFCIFELALTQRKLVKAGQLKPGAQAGGLFGQPA
QNTQGGLFGGGGAATTPGLFGGAQNNTTGGLFGGQNTQAGGGLFGAPNNAAATGLFGAGNANTQGGLFGAKPAATGGGLF
GQPAQTQAGGLFGNTAQPAGGGLFGGATTTPGGGLFGGNTAATGGGLFGGNTQGATGGLFGGQQPNNQGGLFGNTNANTG
GGLFGGATTTTGGGLFGGSTGATGGGLFGGASQPAAGGLFGGAAPQQNSGLFGGATAGQTGGLFGGATQQQGGGLFGQTA
SNPGGGLFGAANATTQPGLFGGNNQAATS

**GLFG//D**$_{52x12}$
**Plasmid: pSNG076**

GLFGNTGDAPAGGLFGNTQDQQGGGLFGQPQDTQQGGLFGQTGDTTNGGLFGGATDTAPGGLFGGGGDNPTGGLFGGNND
QQTGGLFGQGTDQTGGGLFGAPQDNQNGGLFGGGTDTTGGGLFGANTDTQGGGLFGGPSDPTTAGLFGSNNDTTGGGLFG
QPADTNNGGLFGGQTDNQASGLFGANNDPPTNGLFGNNNDPQTAGLFGGATDTGNTGLFGGANDTNGGGLFGNNTDNPTG
GLFGATNDAGGGGLFGGGADTTGGGLFGGGNDQTGGGLFGTANDTTAGGLFGGGNDQPQNGLFGNNNDPATGGLFGQTND
AAPQGLFGGTNDNAASGLFGQKPDSANGVLTKPNEKNLCYAISNGTDFCIFELALTQRKLVKAGQLKPGAQAGGLFGQPA
DNTQGGLFGGGGDATTPGLFGGAQDNTTGGLFGGQNDQAGGGLFGAPNDAAATGLFGAGNDNTQGGLFGAKPDATGGGLF
GQPADTQAGGLFGNTADPAQGGLFGGATDTPGGGLFGGNTDATGGGLFGGNTDQATGGLFGGQQDNNQGGLFGNTNDNTG
GGLFGGATDTTTGGLFGGSTDATGGGLFGGASDPAAGGLFGGAADQQNSGLFGGATDGQTGGLFGGATDQQGGGLFGQTA
DNPGGGLFGAANDTTQPGLFGGNNDAATTS

**GLFG**$_{52x10}$
**Plasmid: pSNG046**

GLFGNTGGAPGLFGTQTQQGGLFGQPQQQGGLFGQTATTGGLFGGTNTAPGLFGGGGGNTGLFGGNNNQTGLFGQGTQTG
GLFGAPQNNGGLFGGTTTTGGLFGNTQTGGGLFGGPSQPTGLFGSNNPTTGLFGQPATNNGLFGQTNNQAGLFGANNQPT
GLFGNNNKPTGLFGGTTTGNGLFGGANNTGGLFGNNTNPTGLFGATNPAGGLFGGGATGGGLFGGGNTTGGLFGTANTTT
GLFGGNTQPQGLFGNNTPATGLFGQTNNAAGLFGGNNNAAGLFGQKPSANGVLTKPNEKNLCYAISNGTDFCIFELALTQ
RKLVKAGQLKPGAQAGLFGQPANTQGLFGGGAATTGLFGGAQNTTGLFGGQNTQAGLFGAPNNAAGLFGAGNANQGLFGA
KPAAGGLFGQPATQAGLFGNTAQPGGLFGGATTTPGLFGGTAATGGLFGGNTQATGLFGGQQPNNGLFGNTNANGGLFGG
TTTTGGLFGGSTGATGLFGGASQAAGLFGGAAPQQGLFGGAAGQTGLFGGATQQQGLFGQTANPGGLFGAANATQGLFGG
NQAATS

**GLFG$_{52x11}$**
**Plasmid: pSNG042**

GLFGNTGGAPGGLFGTQTQQGGGLFGQPQQQGGGLFGQTATTGGGLFGGTNTAPGGLFGGGGGNTGGLFGGNNNQTGGLF
GQGTQTGGGLFGAPQNNGGGLFGGTTTTGGGLFGNTQTGGGGLFGGPSQPTAGLFGSNNPTTGGLFGQPATNNGGLFGQT
NNQASGLFGANNQPTNGLFGNNNKPTAGLFGGTTTGNTGLFGGANNTGGGLFGNNTNPTGGLFGATNPAGGGLFGGGATG
GGGLFGGGNTTGGGLFGTANTTTGGLFGGNTQPQNGLFGNNTPATGGLFGQTNNAAQGLFGGNNNAASGLFGQKPSANGV
LTKPNEKNLCYAISNGTDFCIFELALTQRKLVKAGQLKPGAQAGGLFGQPANTQGGLFGGGAATTPGLFGGAQNTTGGLF
GGQNTQAGGLFGAPNNAATGLFGAGNANQGGLFGAKPAAGGGLFGQPATQAGGLFGNTAQPGGGLFGGATTTPGGLFGGT
AATGGGLFGGNTQATGGLFGGQQPNNGGLFGNTNANGGGLFGGTTTTGGGLFGGSTGATGGLFGGASQAAGGLFGGAAPQ
QSGLFGGAAGQTGGLFGGATQQQGGLFGQTANPGGGLFGAANATQPGLFGGNQAATS

**GLFG$_{52x13}$**
**Plasmid: pSNG041**

GLFGNTGGTAPAGGLFGNTQTQQQGGGLFGQPQQSTQGGGLFGQTGAGTTGGGLFGGATNTTAPGGLFGGGGGNNPTGGL
FGGNNNPQQTGGLFGQGTTGQTGGGLFGAPQNANQGGGLFGGGTTQTTGGGLFGANTQTTGGGGLFGGPSQGPTTAGLFG
SNNPNTTGGGLFGQPANATNNGGLFGGQTNGNQASGLFGANNQQPPTNGLFGNNNKTPQTAGLFGGATTGTGNTGLFGGA
NNATGGGGLFGNNTNTNPTGGLFGATNPNAGGGGLFGGGATATGGGGLFGGGNTPQTGGGLFGTANTGTTAGGLFGGGNT
TQPQNGLFGNNNTNPATGGLFGQTNNGAAPQGLFGGTNNANAASGLFGQKPAGSANGVLTKPNEKNLCYAISNGTDFCIF
ELALTQRKLVKAGQLKPGAQAGGLFGQPAQQNTQGGLFGGGGAGATTPGLFGGAQNQNTTGGLFGGQNTNQAGGGLFGAP
NNAAAATGLFGAGNAGNTQGGLFGAKPAQATGGGLFGQPAQTTQAGGLFGNTAQGPAGGGLFGGATTNTPGGGLFGGNTA
PATGGGLFGGNTQTGATGGLFGGQQPANNQGGLFGNTNANNTGGGLFGGATTQTTGGGLFGGSTGGATGGGLFGGASQTP
AAGGLFGGAAPAQQNSGLFGGATAGGQTGGLFGGATQPQQGGGLFGQTASNNPGGGLFGAANATTTQPGLFGGNNQAAAT
S

**GLFG$_{52x14}$**
**Plasmid: pSNG047**

GLFGNTGGTGAPAGGLFGNTQTQPQQGGGLFGQPQQSNTQGGGLFGQTGAGQTTGGGLFGGATNTTTAPGGLFGGGGGNA
NPTGGLFGGNNNPGQQTGGLFGQGTTGGQTGGGLFGAPQNATNQGGGLFGGGTTQNTTGGGLFGANTQTPTGGGGLFGGP
SQGAPTTAGLFGSNNPNGTTGGGLFGQPANATTNNGGLFGGQTNGGNQASGLFGANNQQGPPTNGLFGNNNKTTPQTAGL
FGGATTGGTGNTGLFGGANNAGTGGGGLFGNNTNTGNPTGGLFGATNPNTAGGGGLFGGGATAGTGGGGLFGGGNTPGQT
GGGLFGTANTGQTTAGGLFGGGNTTTQPQNGLFGNNNTNGPATGGLFGQTNNGGAAPQGLFGGTNNAANAASGLFGQKPA
GGSANGVLTKPNEKNLCYAISNGTDFCIFELALTQRKLVKAGQLKPGAQAGGLFGQPAQQQNTQGGLFGGGGAGGATTPG
LFGGAQNQTNTTGGLFGGQNTNNQAGGGLFGAPNNATAAATGLFGAGNAGGNTQGGLFGAKPAQNATGGGLFGQPAQTAT
QAGGLFGNTAQGTPAGGGLFGGATTNQTPGGGLFGGNTAPGATGGGLFGGNTQTSGATGGLFGGQQPANNNQGGLFGNTN
ANGNTGGGLFGGATTQGTTGGGLFGGSTGGTATGGGLFGGASQTQPAAGGLFGGAAPAGQQNSGLFGGATAGGGQTGGLF
GGATQPNQQGGGLFGQTASNPNPGGGLFGAANATTTTQPGLFGGNNQAGAATS

**GLFG$_{52x15}$**
**Plasmid: pSNG054**

GLFGNTGGTGAAPAGGLFGNTQTQNTQQGGGLFGQPQQSPNTQGGGLFGQTGAGQPTTGGGLFGGATNTTGTAPGGLFGG
GGGNAANPTGGLFGGNNNPGTQQTGGLFGQGTTGGGQTGGGLFGAPQNATQNQGGGLFGGGTTQNNTTGGGLFGANTQTP
ATGGGGLFGGPSQGATPTTAGLFGSNNPNGPTTGGGLFGQPANATNTNNGGLFGGQTNGGGNQASGLFGANNQQGTPPTN
GLFGNNNKTTQPQTAGLFGGATTGGGTGNTGLFGGANNAGATGGGGLFGNNTNTGNNPTGGLFGATNPNTQAGGGGLFGG
GATAGGTGGGGLFGGGNTPGQQTGGGLFGTANTGQGTTAGGLFGGGNTTTAQPQNGLFGNNNTNGGPATGGLFGQTNNGG
NAAPQGLFGGTNNAATNAASGLFGQKPAGGGSANGVLTKPNEKNLCYAISNGTDFCIFELALTQRKLVKAGQLKPGAQAG
GLFGQPAQQQPNTQGGLFGGGGAGGAATTPGLFGGAQNQTNNTTGGLFGGQNTNNTQAGGGLFGAPNNATAAAATGLFGA
GNAGGGNTQGGLFGAKPAQNTATGGGLFGQPAQTAQTQAGGLFGNTAQGTGPAGGGLFGGATTNQATPGGGLFGGNTAPG
NATGGGLFGGNTQTSGGATGGLFGGQQPANTNNQGGLFGNTNANGQNTGGGLFGGATTQGATTGGGLFGGSTGGTGATGG
GLFGGASQTQNPAAGGLFGGAAPAGPQQNSGLFGGATAGGTGQTGGLFGGATQPNGQQGGGLFGQTASNPQNPGGGLFGA
ANATTSTTQPGLFGGNNQAGTAATS

**GAFG$_{52x12}$**
**Plasmid: pSNG072**
GAFGNTGGAPAGGAFGNTQTQQGGGAFGQPQQTQGGGAFGQTGATTGGGAFGGATNTAPGGAFGGGGGNPTGGAFGGNNN
QQTGGAFGQGTTQTGGGAFGAPQNNQGGGAFGGGTTTTGGGAFGANTQTGGGGAFGGPSQPTTAGAFGSNNPTTGGGAFG
QPANTNNGGAFGGQTNNQASGAFGANNQPPTNGAFGNNNKPQTAGAFGGATTTGNTGAFGGANNTGGGGAFGNNTNNPTG
GAFGATNPAGGGGAFGGGATTGGGGAFGGGNTQTGGGAFGTANTTTAGGAFGGGNTQPQNGAFGNNNTPATGGAFGQTNN
AAPQGAFGGTNNNAASGAFGQKPASANGVLTKPNEKNLCYAISNGTDFCIFELALTQRKLVKAGQLKPGAQAGGAFGQPA
QNTQGGAFGGGGAATTPGAFGGAQNNTTGGAFGGQNTQAGGGAFGAPNNAAATGAFGAGNANTQGGAFGAKPAATGGGAF
GQPAQTQAGGAFGNTAQPAGGGAFGGATTTPGGGAFGGNTAATGGGAFGGNTQGATGGAFGGQQPNNQGGAFGNTNANTG
GGAFGGATTTTGGGAFGGSTGATGGGAFGGASQPAAGGAFGGAAPQQNSGAFGGATAGQTGGAFGGATQQQGGGAFGQTA
SNPGGGAFGAANATTQPGAFGGNNQAATTS

**GLLG$_{52x12}$**
**Plasmid: pSNG124**
GLLGNTGGAPAGGLLGNTQTQQGGGLLGQPQQTQGGGLLGQTGATTGGGLLGGATNTAPGGLLGGGGGNPTGGLLGGNNN
QQTGGLLGQGTTQTGGGLLGAPQNNQGGGLLGGGTTTTGGGLLGANTQTGGGGLLGGPSQPTTAGLLGSNNPTTGGGLLG
QPANTNNGGLLGGQTNNQASGLLGANNQPPTNGLLGNNNKPQTAGLLGGATTTGNTGLLGGANNTGGGGLLGNNTNNPTG
GLLGATNPAGGGGLLGGGATTGGGGLLGGGNTQTGGGLLGTANTTTAGGLLGGGNTQPQNGLLGNNNTPATGGLLGQTNN
AAPQGLLGGTNNNAASGLLGQKPASANGVLTKPNEKNLCYAISNGTDFCIFELALTQRKLVKAGQLKPGAQAGGLLGQPA
QNTQGGLLGGGGAATTPGLLGGAQNNTTGGLLGGQNTQAGGGLLGAPNNAAATGLLGAGNANTQGGLLGAKPAATGGGLL
GQPAQTQAGGLLGNTAQPAGGGLLGGATTTPGGGLLGGNTAATGGGLLGGNTQGATGGLLGGQQPNNQGGLLGNTNANTG
GGLLGGATTTTGGGLLGGSTGATGGGLLGGASQPAAGGLLGGAAPQQNSGLLGGATAGQTGGLLGGATQQQGGGLLGQTA
SNPGGGLLGAANATTQPGLLGGNNQAATS

**GLLG//L$_{52x12}$**
**Plasmid: pSNG131**
GLLGNTGLAPAGGLLGNTQLQQGGGLLGQPQLTQGGGLLGQTGLTTGGGLLGGATLTAPGGLLGGGGLNPTGGLLGGNNL
QQTGGLLGQGTLQTGGGLLGAPQLNQGGGLLGGGTLTTGGGLLGANTLTGGGGLLGGPSLPTTAGLLGSNNLTTGGGLLG
QPALTNNGGLLGGQTLNQASGLLGANNLPPTNGLLGNNNLPQTAGLLGGATLTGNTGLLGGANLTGGGGLLGNNTLNPTG
GLLGATNLAGGGGLLGGGALTGGGGLLGGGNLQTGGGLLGTANLTTAGGLLGGGNLQPQNGLLGNNNLPATGGLLGQTNL
AAPQGLLGGTNLNAASGLLGQKPLSANGVLTKPNEKNLCYAISNGTDFCIFELALTQRKLVKAGQLKPGAQAGGLLGQPA
LNTQGGLLGGGGLATTPGLLGGAQLNTTGGLLGGQNLQAGGGLLGAPNLAAATGLLGAGNLNTQGGLLGAKPLATGGGLL
GQPALTQAGGLLGNTALPAGGGLLGGATLTPGGGLLGGNTLATGGGLLGGNTLGATGGLLGGQQLNNQGGLLGNTNLNTG
GGLLGGATLTTGGGLLGGSTLATGGGLLGGASLPAAGGLLGGAALQQNSGLLGGATLGQTGGLLGGATLQQGGGLLGQTA
LNPGGGLLGAANLTTQPGLLGGNNLAATTS

**GXFG//L$_{52x12}$**
**Plasmid: pSNG087**
GGFGNTGLAPAGGTFGNTQLQQGGGQFGQPQLTQGGGAFGQTGLTTGGGNFGGATLTAPGGGFGGGGLNPTGGNFGGNNL
QQTGGTFGQGTLQTGGGNFGAPQLNQGGGTFGGGTLTTGGGQFGANTLTGGGGQFGGPSLPTTAGPFGSNNLTTGGGNFG
QPALTNNGGNFGGQTLNQASGQFGANNLPPTNGKFGNNNLPQTAGTFGGATLTGNTGNFGGANLTGGGGNFGNNTLNPTG
GPFGATNLAGGGGTFGGGALTGGGGTFGGGNLQTGGGTFGTANLTTAGGTFGGGNLQPQNGTFGNNNLPATGGNFGQTNL
AAPQGNFGGTNLNAASGAFGQKPLSANGVLTKPNEKNLCYAISNGTDFCIFELALTQRKLVKAGQLKPGAQAGGQFGQPA
LNTQGGAFGGGGLATTPGNFGGAQLNTTGGTFGGQNLQAGGGNFGAPNLAAATGAFGAGNLNTQGGAFGAKPLATGGGQF
GQPALTQAGGQFGNTALPAGGGTFGGATLTPGGGAFGGNTLATGGGQFGGNTLGATGGPFGGQQLNNQGGAFGNTNLNTG
GGTFGGATLTTGGGGFGGSTLATGGGQFGGASLPAAGGPFGGAALQQNSGAFGGATLGQTGGQFGGATLQQGGGSFGQTA
LNPGGGAFGAANLTTQPGQFGGNNLAATTS

**GFLG$_{52x12}$**
**Plasmid: pSNG081**
GFLGNTGGAPAGGFLGNTQTQQGGGFLGQPQQTQGGGFLGQTGATTGGGFLGGATNTAPGGFLGGGGGNPTGGFLGGNNN
QQTGGFLGQGTTQTGGGFLGAPQNNQGGGFLGGGTTTTGGGFLGANTQTGGGGFLGGPSQPTTAGFLGSNNPTTGGGFLG
QPANTNNGGFLGGQTNNQASGFLGANNQPPTNGFLGNNNKPQTAGFLGGATTTGNTGFLGGANNTGGGGFLGNNTNNPTG
GFLGATNPAGGGGFLGGGATTGGGGFLGGGNTQTGGGFLGTANTTTAGGFLGGGNTQPQNGFLGNNNTPATGGFLGQTNN
AAPQGFLGGTNNNAASGFLGQKPASANGVLTKPNEKNLCYAISNGTDFCIFELALTQRKLVKAGQLKPGAQGGFLGQPAQ
NTQGGFLGGGGAATTPGFLGGAQNNTTGGFLGGQNTQAGGGFLGAPNNAAATGFLGAGNANTQGGFLGAKPAATGGGFLG
QPAQTQAGGFLGNTAQPAGGGFLGGATTTPGGGFLGGNTAATGGGFLGGNTQGATGGFLGGQQPNNQGGFLGNTNANTGG
GFLGGATTTTGGGFLGGSTGATGGGFLGGASQPAAGGFLGGAAPQQNSGFLGGATAGQTGGFLGGATQQQGGGFLGQTAS
NPGGGFLGAANATTQPGFLGGNNQAATS

**GLYG$_{52x12}$**
**Plasmid: pSNG097**
GLYGNTGGAPAGGLYGNTQTQQGGGLYGQPQQTQGGGLYGQTGATTGGGLYGGATNTAPGGLYGGGGGNPTGGLYGGNNN
QQTGGLYGQGTTQTGGGLYGAPQNNQGGGLYGGGTTTTGGGLYGANTQTGGGGLYGGPSQPTTAGLYGSNNPTTGGGLYG
QPANTNNGGLYGGQTNNQASGLYGANNQPPTNGLYGNNNKPQTAGLYGGATTTGNTGLYGGANNTGGGGLYGNNTNNPTG
GLYGATNPAGGGGLYGGGATTGGGGLYGGGNTQTGGGLYGTANTTTAGGLYGGGNTQPQNGLYGNNNTPATGGLYGQTNN
AAPQGLYGGTNNNAASGLYGQKPASANGVLTKPNEKNLCYAISNGTDFCIFELALTQRKLVKAGQLKPGAQAGGLYGQPA
QNTQGGLYGGGGAATTPGLYGGAQNNTTGGLYGGQNTQAGGGLYGAPNNAAATGLYGAGNANTQGGLYGAKPAATGGGLY
GQPAQTQAGGLYGNTAQPAGGGLYGGATTTPGGGLYGGNTAATGGGLYGGNTQGATGGLYGGQQPNNQGGLYGNTNANTG
GGLYGGATTTTGGGLYGGSTGATGGGLYGGASQPAAGGLYGGAAPQQNSGLYGGATAGQTGGLYGGATQQQGGGLYGQTA
SNPGGGLYGAANATTQPGLYGGNNQAATTS

**GAYG$_{52x12}$**
**Plasmid: pSNG098**
GAYGNTGGAPAGGAYGNTQTQQGGGAYGQPQQTQGGGAYGQTGATTGGGAYGGATNTAPGGAYGGGGGNPTGGAYGGNNN
QQTGGAYGQGTTQTGGGAYGAPQNNQGGGAYGGGTTTTGGGAYGANTQTGGGGAYGGPSQPTTAGAYGSNNPTTGGGAYG
QPANTNNGGAYGGQTNNQASGAYGANNQPPTNGAYGNNNKPQTAGAYGGATTTGNTGAYGGANNTGGGGAYGNNTNNPTG
GAYGATNPAGGGGAYGGGATTGGGGAYGGGNTQTGGGAYGTANTTTAGGAYGGGNTQPQNGAYGNNNTPATGGAYGQTNN
AAPQGAYGGTNNNAASGAYGQKPASANGVLTKPNEKNLCYAISNGTDFCIFELALTQRKLVKAGQLKPGAQAGGAYGQPA
QNTQGGAYGGGGAATTPGAYGGAQNNTTGGAYGGQNTQAGGGAYGAPNNAAATGAYGAGNANTQGGAYGAKPAATGGGAY
GQPAQTQAGGAYGNTAQPAGGGAYGGATTTPGGGAYGGNTAATGGGAYGGNTQGATGGAYGGQQPNNQGGAYGNTNANTG
GGAYGGATTTTGGGAYGGSTGATGGGAYGGASQPAAGGAYGGAAPQQNSGAYGGATAGQTGGAYGGATQQQGGGAYGQTA
SNPGGGAYGAANATTQPGAYGGNNQAATTS

**GIFG$_{52x12}$**
**Plasmid: pSNG067**
GIFGNTGGAPAGGIFGNTQTQQGGGIFGQPQQTQGGGIFGQTGATTGGGIFGGATNTAPGGIFGGGGGNPTGGIFGGNNN
QQTGGIFGQGTTQTGGGIFGAPQNNQGGGIFGGGTTTTGGGIFGANTQTGGGGIFGGPSQPTTAGIFGSNNPTTGGGIFG
QPANTNNGGIFGGQTNNQASGIFGANNQPPTNGIFGNNNKPQTAGIFGGATTTGNTGIFGGANNTGGGGIFGNNTNNPTG
GIFGATNPAGGGGIFGGGATTGGGGIFGGGNTQTGGGIFGTANTTTAGGIFGGGNTQPQNGIFGNNNTPATGGIFGQTNN
AAPQGIFGGTNNNAASGIFGQKPASANGVLTKPNEKNLCYAISNGTDFCIFELALTQRKLVKAGQLKPGAQAGGIFGQPA
QNTQGGIFGGGGAATTPGIFGGAQNNTTGGIFGGQNTQAGGGIFGAPNNAAATGIFGAGNANTQGGIFGAKPAATGGGIF
GQPAQTQAGGIFGNTAQPAGGGIFGGATTTPGGGIFGGNTAATGGGIFGGNTQGATGGIFGGQQPNNQGGIFGNTNANTG
GGIFGGATTTTGGGIFGGSTGATGGGIFGGASQPAAGGIFGGAAPQQNSGIFGGATAGQTGGIFGGATQQQGGGIFGQTA
SNPGGGIFGAANATTQPGIFGGNNQAATS

**GLFA$_{52x12}$**
**Plasmid: pSNG068**
GLFANTGGAPAGGLFANTQTQQGGGLFAQPQQTQGGGLFAQTGATTGGGLFAGATNTAPGGLFAGGGGNPTGGLFAGNNN
QQTGGLFAQGTTQTGGGLFAAPQNNQGGGLFAGGTTTTGGGLFAANTQTGGGGLFAGPSQPTTAGLFASNNPTTGGGLFA
QPANTNNGGLFAGQTNNQASGLFAANNQPPTNGLFANNNKPQTAGLFAGATTTGNTGLFAGANNTGGGGLFANNTNNPTG
GLFAATNPAGGGGLFAGGATTGGGGLFAGGNTQTGGGLFATANTTTAGGLFAGGNTQPQNGLFANNNTPATGGLFAQTNN
AAPQGLFAGTNNNAASGLFAQKPASANGVLTKPNEKNLCYAISNGTDFCIFELALTQRKLVKAGQLKPGAQAGGLFAQPA
QNTQGGLFAGGAATTPGLFAGAQNNTTGGLFAGQNTQAGGGLFAAPNNAAATGLFAAGNANTQGGLFAAKPAATGGGLF
AQPAQTQAGGLFANTAQPAGGGLFAGATTTPGGGLFAGNTAATGGGLFAGNTQGATGGLFAGQQPNNQGGLFANTNANTG
GGLFAGATTTTGGGLFAGSTGATGGGLFAGASQPAAGGLFAGAAPQQNSGLFAGATAGQTGGLFAGATQQQGGGLFAQTA
SNPGGGLFAAANATTQPGLFAGNNQAATS

**SLFG$_{52x12}$**
**Plasmid: pSNG073**
SLFGNTGGAPAGSLFGNTQTQQGGSLFGQPQQTQGGSLFGQTGATTGGSLFGGATNTAPGSLFGGGGGNPTGSLFGGNNN
QQTGSLFGQGTTQTGGSLFGAPQNNQGGSLFGGGTTTTGGSLFGANTQTGGGSLFGGPSQPTTASLFGSNNPTTGGSLFG
QPANTNNGSLFGGQTNNQASSLFGANNQPPTNSLFGNNNKPQTASLFGGATTTGNTSLFGGANNTGGGSLFGNNTNNPTG
SLFGATNPAGGGSLFGGGATTGGGSLFGGGNTQTGGSLFGTANTTTAGSLFGGGNTQPQNSLFGNNNTPATGSLFGQTNN
AAPQSLFGGTNNNAASSLFGQKPASANGVLTKPNEKNLCYAISNGTDFCIFELALTQRKLVKAGQLKPGAQAGSLFGQPA
QNTQGSLFGGGGAATTPSLFGGAQNNTTGSLFGGQNTQAGGSLFGAPNNAAATSLFGAGNANTQGSLFGAKPAATGGSLF
GQPAQTQAGSLFGNTAQPAGGSLFGGATTTPGGSLFGGNTAATGGSLFGGNTQGATGSLFGGQQPNNQGSLFGNTNANTG
GSLFGGATTTTGGSLFGGSTGATGGSLFGGASQPAAGSLFGGAAPQQNSSLFGGATAGQTGSLFGGATQQQGGSLFGQTA
SNPGGSLFGAANATTQPSLFGGNNQAATS

**GLFS$_{52x12}$**
**Plasmid: pSNG082**
GLFSNTGGAPAGGLFSNTQTQQGGGLFSQPQQTQGGGLFSQTGATTGGGLFSGATNTAPGGLFSGGGGNPTGGLFSGNNN
QQTGGLFSQGTTQTGGGLFSAPQNNQGGGLFSGGTTTTGGGLFSANTQTGGGGLFSGPSQPTTAGLFSSNNPTTGGGLFS
QPANTNNGGLFSGQTNNQASGLFSANNQPPTNGLFSNNNKPQTAGLFSGATTTGNTGLFSGANNTGGGGLFSNNTNNPTG
GLFSATNPAGGGGLFSGGATTGGGGLFSGGNTQTGGGLFSTANTTTAGGLFSGGNTQPQNGLFSNNNTPATGGLFSQTNN
AAPQGLFSGTNNNAASGLFSQKPASANGVLTKPNEKNLCYAISNGTDFCIFELALTQRKLVKAGQLKPGAQGGLFSQPAQ
NTQGGLFSGGGAATTPGLFSGAQNNTTGGLFSGQNTQAGGGLFSAPNNAAATGLFSAGNANTQGGLFSAKPAATGGGLFS
QPAQTQAGGLFSNTAQPAGGGLFSGATTTPGGGLFSGNTAATGGGLFSGNTQGATGGLFSGQQPNNQGGLFSNTNANTGG
GLFSGATTTTGGGLFSGSTGATGGGLFSGASQPAAGGLFSGAAPQQNSGLFSGATAGQTGGLFSGATQQQGGGLFSQTAS
NPGGGLFSAANATTQPGLFSGNNQAATS

**FSFG$_{52x12}$**
**Plasmid: pSNG069**
FSFGNTGGAPAGFSFGNTQTQQGGFSFGQPQQTQGGFSFGQTGATTGGFSFGGATNTAPGFSFGGGGGNPTGFSFGGNNN
QQTGFSFGQGTTQTGGFSFGAPQNNQGGFSFGGGTTTTGGFSFGANTQTGGGFSFGGPSQPTTAFSFGSNNPTTGGFSFG
QPANTNNGFSFGGQTNNQASFSFGANNQPPTNFSFGNNNKPQTAFSFGGATTTGNTFSFGGANNTGGGFSFGNNTNNPTG
FSFGATNPAGGGFSFGGGATTGGGFSFGGGNTQTGGFSFGTANTTTAGFSFGGGNTQPQNFSFGNNNTPATGFSFGQTNN
AAPQFSFGGTNNNAASFSFGQKPASANGVLTKPNEKNLCYAISNGTDFCIFELALTQRKLVKAGQLKPGAQAGFSFGQPA
QNTQGFSFGGGGAATTPFSFGGAQNNTTGFSFGGQNTQAGGFSFGAPNNAAATFSFGAGNANTQGFSFGAKPAATGGFSF
GQPAQTQAGFSFGNTAQPAGGFSFGGATTTPGGFSFGGNTAATGGFSFGGNTQGATGFSFGGQQPNNQGFSFGNTNANTG
GFSFGGATTTTGGFSFGGSTGATGGFSFGGASQPAAGFSFGGAAPQQNSFSFGGATAGQTGFSFGGATQQQGGFSFGQTA
SNPGGFSFGAANATTQPFSFGGNNQAATS

**FSFG$_{52x12}$ F$\rightarrow$S mutant 1**
**Plasmid: pSNG150**
FSFGNTGGAPAGFSFGNTQTQQGGSSSGQPQQTQGGFSFGQTGATTGGFSFGGATNTAPGSSSGGGGGNPTGFSFGGNNN
QQTGFSFGQGTTQTGGSSSGAPQNNQGGFSFGGGTTTTGGFSFGANTQTGGGSSSGGPSQPTTAFSFGSNNPTTGGFSFG
QPANTNNGSSSGGQTNNQASFSFGANNQPPTNFSFGNNNKPQTASSSGGATTTGNTFSFGGANNTGGGFSFGNNTNNPTG
SSSGATNPAGGGFSFGGGATTGGGFSFGGGNTQTGGSSSGTANTTTAGFSFGGGNTQPQNFSFGNNNTPATGSSSGQTNN
AAPQFSFGGTNNNAASFSFGQKPASANGVLTKPNEKNLCYAISNGTDFCIFELALTQRKLVKAGQLKPGAQAGSSSGQPA
QNTQGFSFGGGGAATTPFSFGGAQNNTTGSSSGGQNTQAGGFSFGAPNNAAATFSFGAGNANTQGSSSGAKPAATGGFSF
GQPAQTQAGFSFGNTAQPAGGSSSGGATTTPGGFSFGGNTAATGGFSFGGNTQGATGSSSGGQQPNNQGFSFGNTNANTG
GFSFGGATTTTGGSSSGGSTGATGGFSFGGASQPAAGFSFGGAAPQQNSSSSGGATAGQTGFSFGGATQQQGGFSFGQTA
SNPGGSSSGAANATTQPFSFGGNNQAATS

**FSFG$_{52x12}$ F$\rightarrow$S mutant 2**
**Plasmid: pSNG151**
FSFGNTGGAPAGFSFGNTQTQQGGFSFGQPQQTQGGFSFGQTGATTGGSSSGGATNTAPGSSSGGGGGNPTGFSFGGNNN
QQTGFSFGQGTTQTGGFSFGAPQNNQGGFSFGGGTTTTGGSSSGANTQTGGGSSSGGPSQPTTAFSFGSNNPTTGGFSFG
QPANTNNGFSFGGQTNNQASFSFGANNQPPTNSSSGNNNKPQTASSSGGATTTGNTFSFGGANNTGGGFSFGNNTNNPTG
FSFGATNPAGGGFSFGGGATTGGGSSSGGGNTQTGGSSSGTANTTTAGFSFGGGNTQPQNFSFGNNNTPATGFSFGQTNN
AAPQFSFGGTNNNAASSSSGQKPASANGVLTKPNEKNLCYAISNGTDFCIFELALTQRKLVKAGQLKPGAQAGSSSGQPA
QNTQGFSFGGGGAATTPFSFGGAQNNTTGFSFGGQNTQAGGFSFGAPNNAAATSSSGAGNANTQGSSSGAKPAATGGFSF
GQPAQTQAGFSFGNTAQPAGGFSFGGATTTPGGFSFGGNTAATGGSSSGGNTQGATGSSSGGQQPNNQGFSFGNTNANTG
GFSFGGATTTTGGFSFGGSTGATGGFSFGGASQPAAGSSSGGAAPQQNSSSSGGATAGQTGFSFGGATQQQGGFSFGQTA
SNPGGFSFGAANATTQPFSFGGNNQAATS

**GLFG$_{29x12}$**
**Plasmid: pSNG148**
GLFGNTGGAPAGGLFGNTQTQQGGGLFGQPQQTQGGGLFGQTGATTGGGLFGGATNTAPGGLFGGGGGNPTGGLFGGNNN
QQTGGLFGQGTTQTGGGLFGAPQNNQGGGLFGGGTTTTGGGLFGANTQTGGGGLFGGPSQPTTAGLFGSNNPTTGGGLFG
QPANTNNGGLFGGQTNNQASGLFGANNQPPTNGLFGNNNKPQTAGLFGGATTTGNTGLFGGANNTGGGGLFGNNTNNPTG
GLFGATNPAGGGGLFGGGATTGGGGLFGGGNTQTGGGLFGTANTTTAGGLFGGGNTQPQNGLFGNNNTPATGGLFGQTNN
AAPQGLFGGTNNNAASGLFGQKPASANGVLT

**FSFG$_{29x12}$**
**Plasmid: pSNG147**
FSFGNTGGAPAGFSFGNTQTQQGGFSFGQPQQTQGGFSFGQTGATTGGFSFGGATNTAPGFSFGGGGGNPTGFSFGGNNN
QQTGFSFGQGTTQTGGFSFGAPQNNQGGFSFGGGTTTTGGFSFGANTQTGGGFSFGGPSQPTTAFSFGSNNPTTGGFSFG
QPANTNNGFSFGGQTNNQASFSFGANNQPPTNFSFGNNNKPQTAFSFGGATTTGNTFSFGGANNTGGGFSFGNNTNNPTG
FSFGATNPAGGGFSFGGGATTGGGFSFGGGNTQTGGFSFGTANTTTAGFSFGGGNTQPQNFSFGNNNTPATGFSFGQTNN
AAPQFSFGGTNNNAASFSFGQKPASANGVLT

**FSFG$_{52x15}$**
**Plasmid: pSNG080**

FSFGNTGGTGAAPAGFSFGNTQTQNTQQGGFSFGQPQQSPNTQGGFSFGQTGAGQPTTGGFSFGGATNTTGTAPGFSFGG
GGGNAANPTGFSFGGNNNPGTQQTGFSFGQGTTGGGQTGGFSFGAPQNATQNQGGFSFGGGTTQNNTTGGFSFGANTQTP
ATGGGFSFGGPSQGATPTTAFSFGSNNPNGPTTGGFSFGQPANATNTNNGFSFGGQTNGGGNQASFSFGANNQQGTPPTN
FSFGNNNKTTQPQTAFSFGGATTGGGTGNTFSFGGANNAGATGGGFSFGNNTNTGNNPTGFSFGATNPNTQAGGGFSFGG
GATAGGTGGGFSFGGGNTPGQQTGGFSFGTANTGQGTTAGFSFGGGNTTTAQPQNFSFGNNNTNGGPATGFSFGQTNNGG
NAAPQFSFGGTNNAATNAASFSFGQKPAGGGSANGVLTKPNEKNLCYAISNGTDFCIFELALTQRKLVKAGQLKPGAQGF
SFGQPAQQQPNTQGFSFGGGGAGGAATTPFSFGGAQNQTNNTTGFSFGGQNTNNTQAGGFSFGAPNNATAAAATFSFGAG
NAGGGNTQGFSFGAKPAQNTATGGFSFGQPAQTAQTQAGFSFGNTAQGTGPAGGFSFGGATTNQATPGGFSFGGNTAPGN
ATGGFSFGGNTQTSGGATGFSFGGQQPANTNNQGFSFGNTNANGQNTGGFSFGGATTQGATTGGFSFGGSTGGTGATGGF
SFGGASQTQNPAAGFSFGGAAPAGPQQNSFSFGGATAGGTGQTGFSFGGATQPNGQQGGFSFGQTASNPQNPGGFSFGAA
NATTSTTQPFSFGGNNQAGTAATS

**FSFG$_{52x18}$**
**Plasmid: pSNG086**

FSFGNTGGTGAAGTAPAGFSFGNTQTQNTNPQQQGGFSFGQNQQSPNTTSTQGGFSFGQTGAGQPGGTTTGGFSFGGATN
TTGTQNTAPGFSFGGGGGNAAGTGNPTGFSFGGNNNPGTQGAQQTGFSFGQGTTSGGPNGQTGGFSFGAPQNATQNGQNQ
GGFSFGGGTTQNNAGPTTGGFSFGANTQTPAANTTGGGFSFGGPSQGATTSGGTTAFSFGSNNPNGNAAGTTGGFSFGQP
ANATNGTGTNNGFSFGGQTNAGGPGNNQASFSFGANNQQGTTAQAPTNFSFGNNNKTTQQNTPQTAFSFGGATTGGAQQP
TGNTFSFGGANNAGAPGQTGGGFSFGNNTNTGNPGGNPTGFSFGATNPNTQATAAGGGFSFGGGATAGGNGPTGGGFSFG
GGNTPGQNNTQTGGFSFGTANTGQGTPATTAGFSFGGGNTTTAQTNQPQNFSFGNNNTNGGTGAPATGFSFGQTNNGGNQ
QNAAPQFSFGGTNNAATPGTNAASFSFGQKPAGGNAATSANGVLTKPNEKNLCYAISNGTDFCIFELALTQRKLVKAGQL
KPGAQAGFSFGQPAQQQGGNNNTQGFSFGGGGAGGANPGGATTFSFGGAQNQTNGQPNTTGFSFGGQNTNNTSPNQAGGF
SFGAPNNATATTAAAATFSFGAGNAGNGAGGPTQGFSFGAKPAQNTGQGATGGFSFGQPAQTAQNTQTQAGFSFGNTAQG
TATGNPAGGFSFGGATTNQAAGATPGGFSFGGNTAPGNGAGATGGFSFGGNTQTSGTTQPATGFSFGGQQPANTQGTNNQ
GFSFGNTNANGQAGGPTGGFSFGGATTQGAATNTTPGFSFGGSTGGTGGATPTGGFSFGGASQTQNGPGGAAGFSFGGAA
PAGQNNGQQNSFSFGGATAGGTPATQQTGFSFGGATQPNGGPGQQGGFSFGQTASNPQNAANAGGFSFGAANATTSPTTG
TTQFSFGGNNQAGTQNTPATS
